# Early *C. elegans* embryos modulate cell division timing to compensate for, and survive, the discordant conditions of a severe temperature gradient

**DOI:** 10.1101/2020.06.02.128694

**Authors:** Eric Terry, Bilge Birsoy, David Bothman, Marin Sigurdson, Pradeep M. Joshi, Carl Meinhart, Joel H. Rothman

## Abstract

Despite a constant barrage of intrinsic and environmental noise, embryogenesis is remarkably reliable, suggesting the existence of systems that ensure faithful execution of this complex process. We report that early *C. elegans* embryos, which normally show a highly reproducible lineage and cellular geometry, can compensate for deviations imposed by the discordant conditions of a steep temperature gradient generated in a microfluidic device starting at the two-cell stage. Embryos can survive a gradient of up to 7.5°C across the 50-micron axis through at least three rounds of division. This response is orientation-dependent: survival is higher when the normally faster-dividing anterior daughter of the zygote, AB, but not its sister, the posterior P1, is warmer. We find that temperature-dependent cellular division rates in the early embryo can be effectively modeled by a modification of the Arrhenius equation. Further, both cells respond to the gradient by dramatically reducing division rates compared to the predicted rates for the temperature experienced by the cell even though the temperature extremes are well within the range for normal development. This finding suggests that embryos may sense discordance and slow development in response. We found that in the cohort of surviving embryos, the cell on the warmer side at the two-cell stage shows a greater average decrease in expected division rate than that on the cooler side, thereby preserving the normal cellular geometry of the embryo under the discordant conditions. A diminished average slow-down response correlated with lethality, presumably owing to disruption of normal division order and developmental fidelity. Remarkably, some inviable embryos in which the canonical division order was reversed nonetheless proceeded through relatively normal morphogenesis, suggesting a subsequent compensation mechanism independent of cell division control. These findings provide evidence for a previously unrecognized process in *C. elegans* embryos that may serve to compensate for deviations imposed by aberrant environmental conditions, thereby resulting in a high-fidelity output.

## Introduction

The development of a complex multicellular animal from a zygote requires coordination of diverse biological processes. Each step in the process is associated with a particular rate of error and is subject to perturbation by genetic variation, environmental fluctuations, and intrinsic molecular noise [1]. Nonetheless, despite the incessant onslaught of error-provoking influences, development generally proceeds faithfully in organisms spanning metazoan phylogeny [2–8]. The observed rate of success is remarkable: throughout the complex process of embryogenesis, cells must properly satisfy many parameters of identity and behavior, including appropriate gene expression, incidence and timing of cell division, and spatiotemporal positioning. This ability of development to proceed with high fidelity in the face of environmental and intrinsic variation likely reflects evolutionary selection for robustness-conferring cellular and molecular mechanisms [1].

While the processes that regulate spatiotemporal developmental fidelity have not been comprehensively elucidated, several mechanisms that influence developmental precision have been uncovered. The molecular chaperone, Hsp90, has been found to act as a buffer against cryptic variation in both fly and vertebrate models [9–14], helping to ensure appropriate cellular identity during development. Robustness in spatial coordination is exemplified by transformation of a variable Bicoid gradient along the antero-posterior axis of the *Drosophila* embryo into spatially stereotyped expression pattern of hunchback and ultimately cellular identity along the anterior-posterior axis of developing embryos [3,15,16]. In vertebrates, Notch directs intercellular temporal coordination and precision during somite development [17] and mutations in the Notch signaling pathway result in the loss of the normal synchrony in division oscillations of somite cells [17–19]. While such findings shed light on processes that regulate developmental precision, systems that mediate developmental fidelity, particularly in the temporal dimension, are not well understood.

The early cell divisions in *C. elegans* embryos establish six “founder cell” lineages. While the founder cells are born by a series of asynchronous divisions, all their descendants divide in approximate synchrony with a cell cycle periodicity that is unique to that lineage [20]. Although the different founder cell division clocks are not synchronized with each other, they must be kept in precise register to ensure the highly stereotyped arrangement of cells that is characteristic of early embryogenesis [20,21]. This reproducible geometry is critical for signaling events that depend on the proper spatial arrangements of cell-cell contacts that are essential for normal development to proceed [22,23].

The differential apportionment of cell division clocks is evident as early as the two-cell stage, in which the larger, anterior daughter of the zygote, the AB blastomere, divides before its posterior sister, P1, with high precision. This difference in cell cycle timing is passed onto the descendants of each cell, such that the cell cycle periodicity in the AB lineage is shorter than in the P_1_ lineage. The regulation of this differential timing control system has been thought to be determined by cell-autonomous mechanisms; when isolated AB and P_1_ are allowed to develop in culture, the relative division timing difference is largely preserved [20,24–28]. The difference in the cell cycle clocks of AB and P_1_ largely correlates with their different molecular content and size [25,29,30], which are controlled by the machinery that establishes anteroposterior polarity in the zygote following fertilization [31–33].

The high fidelity of molecular processes such as DNA replication and aminoacyl-tRNA charging [34,35] result from reactions that correct errors after they are made and it is conceivable that errors made in embryogenesis are often corrected through subsequent developmental processes. Indeed, development is characterized by substantial plasticity and regulatory mechanisms often generate precise patterning from more disordered assemblages of cells (e.g. ordered patterning of hair follicles [36] and robust patterning by morphogen gradients [37–43]). When cells in different domains of *Drosophila* embryos are forced to develop at different rates by imposition of a temperature (T) gradient, abnormal patterns of cell divisions arise (with fewer nuclei on the colder side); however, developmental gene expression is resolved into normal patterns along the axis [44,45], revealing that the gene expression patterning machinery can correct for abnormal cellular patterns that were induced by discordant conditions occurring earlier in development.

The highly stereotypical division sequence and arrangement of early blastomeres in the rapidly developing *C. elegans* embryo [20,21,24,46,47] persists across developmental rates that vary over nearly an order of magnitude (dependent on the T of the environment), providing a useful system for probing mechanisms that ensure developmental precision and fidelity. How are the founder cell lineages, each with different cell cycle clocks, coordinated irrespective of developmental rate, environmental variation, and intrinsic molecular noise, ensuring a reproducible outcome? It is conceivable that communication between cells in different lineages functions to tune cell division timings while they are occurring, thereby continuously maintaining proper harmony across the developing embryo. Alternatively, deviations might be compensated by subsequent error-correcting responses that renormalizes cellular geometry.

If early *C. elegans* embryos harbor mechanisms that correct for environmental variation, then they may undergo stereotypical development even under the discordant conditions imposed by a T gradient that, in the absence of such correction, would drive cell divisions and placements to deviate from the normal pattern.

In this study, we investigated whether developing *C. elegans* embryos can compensate and correct for discordant conditions between lineages by subjecting them to steep T gradients along the long (anteroposterior) axis. To achieve this, we designed, fabricated, and validated a novel microfluidic device that establishes a steep T gradient in which the two extremes are nonetheless within the permissive T ranges for normal development. While embryos in this steep gradient would be predicted to undergo out-of-sequence division patterns and die in the absence of correction, remarkably, we found that embryos can survive a T gradient of up to 7.5°C across the 50-micron axis through at least three rounds of division from first cleavage, suggesting that they can compensate for the large discordance imposed by the gradient. This response showed orientation-dependence: survival was higher when the normally faster-dividing anterior daughter of the zygote, AB, but not its posterior sister, P_1_, was at the warmer T. We found that the division timing of both AB and P_1_ slowed down dramatically in the presence of the gradient compared to the predicted rates, suggesting that embryos may sense and respond to the “crisis” of a discordant condition by activating a checkpoint-like system. Further, cells on the warmer side slow by a larger extent relative to predicted rates than cells on the cooler side, with the result that the normal division sequence and geometry are preserved. The magnitude of this response correlated with embryo survival: those with a stronger “tuning” response (adjustment in division rate) showed a higher tendency to survive, whereas those with a modest response generally died, suggesting that the response ensures normal developmental progression. In the largest T gradient, the canonical division sequence of many embryos was reversed, with a warmer P_1_ dividing ahead of AB. Although such embryos invariably died, some nonetheless showed signs of relatively normal morphogenesis, suggesting a later compensation mechanism can act independently of cell division control. These findings are consistent with the possibility that early *C. elegans* embryos possess mechanisms that sense and correct for noise-induced variation at the time it occurs and respond by adjusting cell division rates to restore the normal pattern of development.

## Results

### Development and validation of a microfluidics T gradient device

We sought to investigate whether early *C. elegans* embryos are capable of responding to, and correcting for, severely discordant environmental conditions by subjecting them to a steep T gradient across their long axis. We posit that if no compensation system exists, this environmental discordance would drive opposite ends of the embryo to develop at different rates. Based on the known relationship between development timing and T[48], a gradient of 5°C across the 50 μm anteroposterior axis would be expected to create an ∼1.5x difference in the developmental rates of P_1_ and AB in the absence of any adjustments made by the embryo. To perform this test for developmental compensation, we designed and fabricated a microfluidic device (Fig. 1; Movie S1; see Materials and Methods) that establishes a gradient of up to 7.5°C across the long axis of the developing embryo, while ensuring that the cells within the embryo remained within permissive, non-stressful Ts for normal development (between 16°C and 24°C).

**Figure 1.**
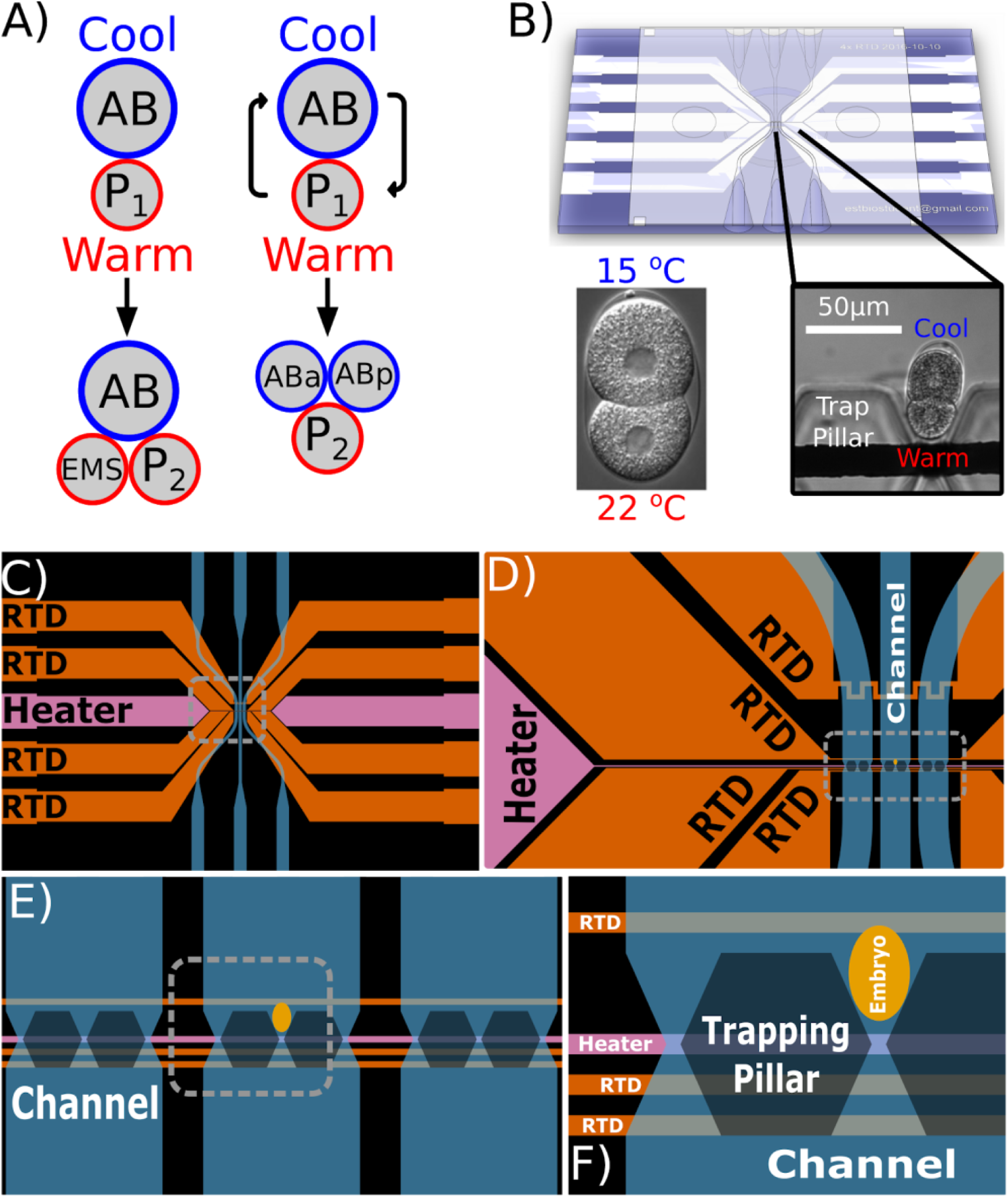
Microfluidic device design. A) The two-cell *C. elegans* embryo is subjected to a T gradient. Normally, in absence of a gradient under constant uniform Ts, AB divides before P_1_. In a gradient AB and P_1_ divide with rates of division determined by the T experienced by the respective cell. In absence of a coordination mechanism between the two cells (left), it is possible to establish a T differential at which the order of division of AB and P_1_ is reversed wherein the warmer P_1_ cell divides before the cooler AB cell. If there were a coordination mechanism (right) that corrects for the discordant conditions, the embryo should resist the T-dependent rates of division allowing for the canonical order of AB and P_1_ divisions. B) Microfluidic device used to capture and orient embryo in a T gradient. In the example shown here the embryo is oriented such that the posterior smaller P_1_ blastomere is closer to the heating element and experiences a warmer T while the larger anterior AB blastomere experiences a cooler T. C-F) Schematic of the layout of device at four scales. C) Macro view of the device. Blue indicates channels, orange indicates T sensors (RTD) and magenta indicates a Joule heater. D) Closer view of T sensing regions of RTDs and capture region. E) View showing all three channels, capture regions for embryos in each channel, Joule heater, and RTDs close to Joule heater. F) Closeup of a single capture region and example of embryo size and placement. Spacing between heater and lower RTDs is 10µm. Width of the heater and RTDs is 10µm

The microfabricated device that generated the T gradient used a platinum Joule micro-heater to establish the high T side of the gradient and a chilled fluid mixture to cool the surface opposite from that containing the heater. The magnitude of the T gradient was controlled by varying the T of the cooling fluid and the power through the Joule heater. Embryos were flowed into the device through microchannels with a syringe pump and properly oriented and trapped between pillars in the microfluidic device (Fig. 1E, F; Movie S2).

Computational model and numerical simulations predicted that the device would effectively generate the desired magnitude of T gradient. We experimentally validated the actual T profile of the device by filling the microchannels through which embryos were delivered with a T-dependent fluorophore, dextran-conjugated rhodamine B (DCRB), and measuring the relative quantum yield (Fig. 2B, C). The measured T profile closely matched that of the modeled distribution (Fig. 2B, D). We modeled heat transfer through the eggshell and cytoplasm to assess whether the T gradient experienced by the embryo is likely to diverge substantially from the measured environment in the channel (see Materials and Methods). Using even extremely conservative parameters, neither the eggshell, nor fluid convection, reduce the T-gradient in the embryo by more than a few percent, hence the device effectively generates a pole-to-pole difference in T experienced by the embryo of up to 7.5°C.

**Figure 2.**
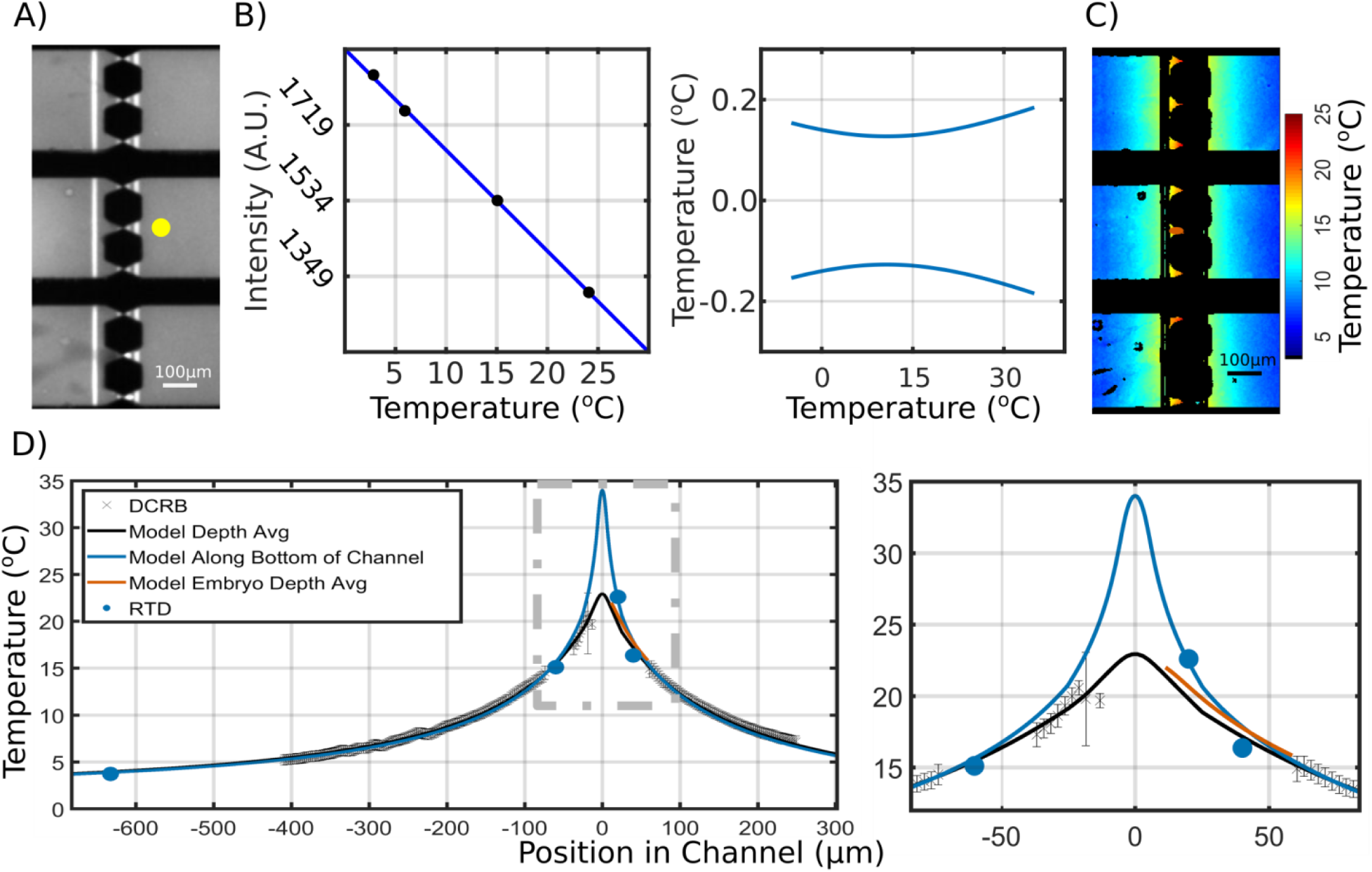
Characterization of the T gradient in the microfluidic device. Characterization and modeling of the T gradient in the microfluidic device using Dextran conjugated Rhodamine B (DCRB). A) Yellow circle represents the pixel intensities of DCRB analyzed. B) (Left) Linear model relating fluorescence intensity of DCRB to T and (right) 97.5% confidence interval distance from model for the inverse linear model of intensity to T, as function of T. C) False coloring heat map of T distribution in device during operation. D) Black data points and error bars indicate the average Ts and standard error across all three channels as a function of position in the channel. X=0 indicates the center of the Joule heating element. Blue line is the model estimate of Ts along the interior bottom of the microfluidic device. Black line is the model estimate of the depth average T in the channel and Orange line is the model estimate of T in the embryo. Blue dots correspond to RTD T measurements at the position relative to the heater

### Embryonic survival is dependent on orientation and magnitude of the T gradient

*C. elegans* embryos develop successfully over a broad T range of 6°C to 26°C. At uniform T, the rate of development and cell division increases by ∼50% for every 5°C increase in T [48]. If cells in a T gradient divide at the rate predicted from the average T experienced and the embryo is unable to correct for this discordance, the division sequence would be expected to diverge substantially from the normal pattern. At the two-cell stage, for example, this could reverse the normal division order, in which the AB cell divides before the P_1_ cell. Such out-of-sequence divisions would be expected to result in aberrant arrangements of cells that diverge substantially from the normally highly stereotyped pattern. Given the rapid intercellular signaling events in the early embryo that are essential for normal cell specification and position [22,29,32,33,49,50] and that depend upon precise alignment of signaling and receiving cells, such derangement of the pattern would be expected to lead to defective embryogenesis, as evidenced by a failure to hatch. Thus, we asked whether discordance imposed by the steep T gradient results in aberrant cell division and geometry and consequent lethality.

We performed an initial assessment to determine whether 1-4-cell stage embryos can survive in a T gradient by loading them into an early version of the device in which the magnitude of the T gradient differed depending on the position of the captured embryo in the device (Fig. 3). This approach allowed us to evaluate hatching as a function of different gradient magnitudes without changing experimental parameters. A flow rate of at least 25 nl/min was essential for adequate oxygen and CO_2_ exchange required to keep embryos viable in the microfluidic device even in absence of a gradient (Fig. 3A). The embryos were subjected to an optimal flow rate of 500 nl/min, which is 20x the critical flow rate necessary for viability without altering the T gradient profile of the microfluidic device (see supplemental text). Early embryos were subjected to the T gradient for ∼1 hour, unloaded from the device, allowed to develop, and scored the following day for hatching on culture plates. We found that the embryos were frequently able to survive through to hatching into viable L1 larvae after exposure to the gradient during the crucial early periods of development. The survival (hatching) rate showed an approximately monotonic decrease with increasing magnitude of the gradient (Fig. 3B). While all embryos survived exposure to a 2°C gradient (n=9), ∼50% survived in a pole-to-pole T gradient of 2.5-3°C (n=20) and ∼25% survived as the magnitude of the gradient was increased from 3°C to 5.5°C (n=46). Under these conditions, none of the embryos exposed to a 6°C T differential hatched (n=5). These initial observations revealed that a) early exposure of *C. elegans* embryos to a T gradient results in significant lethality, b) the degree of lethality correlates with the magnitude of the gradient, and c) some embryos can survive even in very substantial gradient of ∼5.5°C along the long axis.

**Figure 3.**
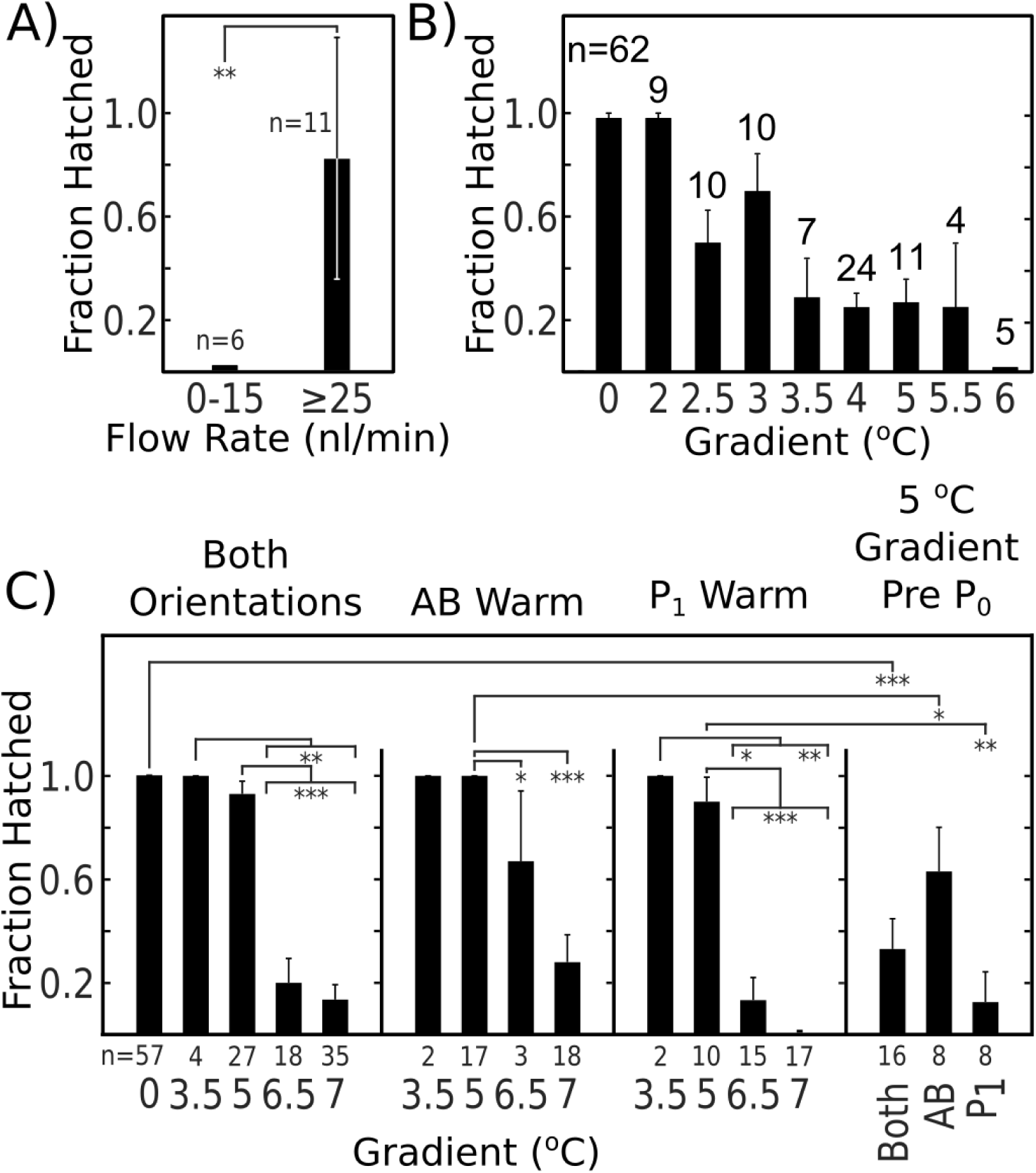
Survival in the T gradient is dependent on both the magnitude of the gradient and orientation of the embryo. A) Fraction of embryos completing development in the microfluidic channel. B) Survival of mixed stage early embryos (1-4 cells) after ∼ 1 hour in the device and then unloaded and placed on agar plates to complete development. C) Survival of embryos in gradients of different magnitudes. Pooled data for embryos in both orientations, embryos with anterior (AB) warm, and embryos with posterior (P_1_) warm in T gradients of different magnitudes are shown. The rightmost graph represents embryos loaded prior to first cell division in a 5°C T gradient (Fisher exact test p<0.05 **p<0.01 ***p<0.001)

To observe individual divisions and characterize the survival of one and two-cell embryos subjected to the gradient as a function of number of divisions, we loaded embryos into a modified device before the division of the first cell, and after the division of the first cell. This allowed us to measure the times of division for each cell while in the gradient. Embryos were loaded into the device in either orientation such that, for some, AB was on the warmer end of the gradient (positioned toward the heater), and for others, P_1_ was warmer. Embryos were allowed to develop in the gradient through the division of the daughters of AB and P_1_, unloaded, allowed to develop at constant T, and scored for hatching ∼24 hours later. Confirming the results with the earlier device, we found that embryos were frequently able to survive a large T gradient (Fig. 3C). The hatching rate of embryos subjected to the gradient starting at the two-cell stage again correlated roughly monotonically with magnitude of the T gradient. Remarkably, we found that some embryos were able survive a very steep pole-to-pole gradient of 7°C.

We observed that the ability of early embryos to survive exposure to the gradient was orientation-dependent. Embryos that were placed in the gradient such that AB was warmer than P_1_ showed a statistically significantly higher (65.8%; n=38) rate of hatching when compared to embryos with the opposite orientation (P_1_ warmer than AB) (28.6% n=42; p=0.0015 Fisher exact test). Above the threshold of a 5°C gradient, survivability in both orientations started to drop, with P_1_ warmer embryos experiencing a significantly larger drop in survival (Fig. 3C). At the highest magnitude of gradient (7°C), while nearly a quarter of the embryos survived when AB was oriented toward the warm end of the gradient (n=18), all embryos (n=17) in the reverse orientation died, a significant difference (p=0.045). These results raise the possibility that the more rapidly dividing AB cell can more effectively “tune” its division rate in response to discordance than the slower dividing P_1_ cell, consistent with our observations of cell cycle timing adjustment (see below).

### Early rates of division described with a modified Arrhenius equation

We sought to test the possibility that the ability of embryos to survive a T gradient might reflect a system that monitors deviations in the early cell division and then adjusts cell division timings to normalize these deviations. To do so, it was necessary to quantify the T-dependent behavior of the cells at uniform constant Ts, and assess whether cells divide at rates that differ from those predicted for their T environment under the discordant conditions of the gradient. If the two-cell *C. elegans* embryo adjusts cell division timings based on this discordance, this effect would be revealed as a tendency for one or both of the cells to divide at a rate other than that expected for the T experienced by that cell. To reveal any such an effect, it was necessary to measure the division rates for AB and P_1_ at a range of constant Ts, and build a quantitative mathematical model of the division time for the second and third divisions in the AB and P_1_ lineages as a function of T (Fig. 4A & B).

**Figure 4.**
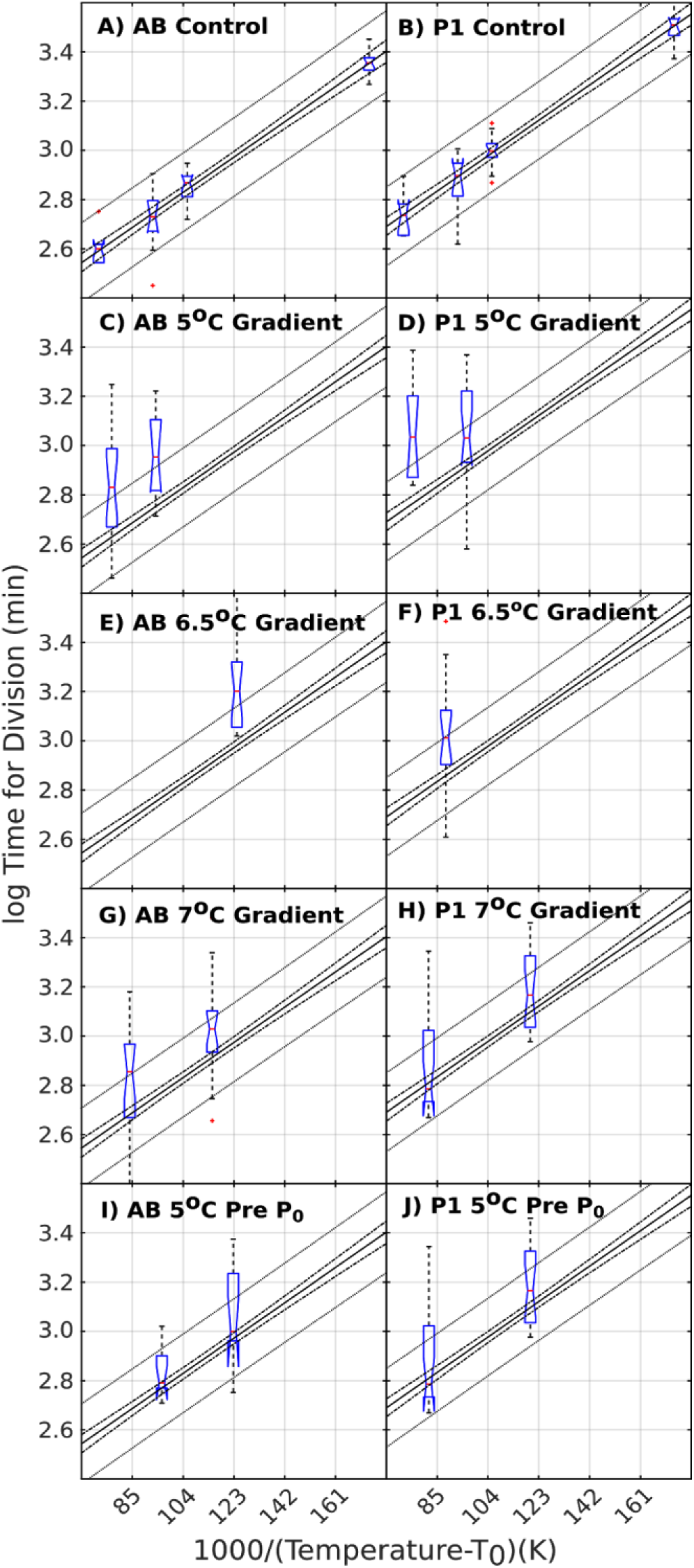
Cell division rates of individual blastomeres in the T gradient. Linear model of division time of AB (A) and P_1_ (B) as a function of T. Notched box plots are data at various Ts. Innermost line indicates linear model. Next outer pair of lines indicate 95% confidence interval while the outermost pair of lines indicate 95% prediction interval. C-H) Notched box plot of division time for AB (left) and P_1_ (right) at the corresponding T plotted over the corresponding model. In both AB and P_1_ plots, the left box corresponds to when the cell is close to the heater and the right box corresponds to when the cell is away from the heater. This cohort of embryos were loaded after formation of first cleavage. I and J) same as C-H except these embryos were loaded in a 5°C gradient before the first cleavage.

We found that the T-dependent times of division for both cells in the two-cell embryo measured at progressively increasing constant T are empirically closely described by a modified Arrhenius equation. This finding is consistent with those described by Begasse et al. [51] for T-dependent rates of pre-division events observed in the one-cell P0 zygote in both *C. elegans* and *C. briggsae*. In that study, as in ours, the data was modeled by performing a least-squares fit to a linearized version of the Arrhenius equation in which the log of the rate (or time interval) of an event is evaluated as a function of the reciprocal of T at which the rate (or time to division) was measured. However, and significantly, our model differs from the previous work in at least one important aspect. The earlier work [51] used the absolute T scale of Kelvin to describe the relationship between rate and T. While this is consistent with calculation of T-dependent chemical rates using the Arrhenius equation, it makes the assumption that the event under consideration progresses at some rate down to a T approaching absolute zero. However, such an assumption does not hold for typical biological processes. In an alternative method introduced by Nakamura et al. [52], an additional T term is introduced into the denominator of the independent variable of the linear form of the Arrhenius equation, allowing for greater empirical fitting of data:

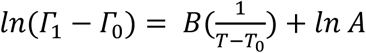

This additional term, which acts as an offset for the measured T of the data, can be thought of as the T at which of the rate for the system under consideration extrapolates to zero. We sought an estimate for this T for *C. elegans* by performing two methods of analysis on our data: a numerical-simulation-generated general non-linear fit of the data, performed in Comsol Multiphysics, and a parametric sweep of this offset T on the linear model of the data. Both methods were in high agreement (∼one part in 100 difference) and revealed that the offset T that best fits our data for the N2 strain of *C. elegans* for the second and third division of the embryo is −10°C. This parameter implies a *C. elegans-* specific “absolute zero” T, at which all cellular activity stops, a more biologically relevant assumption.

### Embryos respond to a T gradient by slowing overall developmental rate

Our modified Arrhenius equation allowed us to calculate an expected rate of cell division at any T within the experimental T ranges and to assess whether the individual cells divided at a rate consistent with, or deviating from, the local T that they experience in the gradient. We followed cell division microscopically throughout exposure to the gradient and quantified the temporal division behavior of each of the cells in the two-cell embryo. This analysis revealed two striking trends in the quantitative behavior of the individual cells within the gradient (Fig. 4C-H).

First, we found that the timing of cell divisions in the gradient showed much greater variability than that observed for embryos at constant T. To compare the variance of cohorts of embryos across the various conditions, we calculated the coefficient of variation (CV) for each cohort of embryos at each of the constant Ts, as well as the CV of each cohort of embryos that experienced the same T gradient magnitude and orientation. The mean coefficient of variation across constant Ts for both AB and P_1_ were 0.10, and the standard deviation of the CVs across the different constant Ts were 0.05 and 0.04 respectively. The mean CV of the various cohorts of embryos experiencing the T gradient was 0.19 and 0.18 for AB and P_1_ respectively with a standard deviation of CVs across gradients and orientations of 0.06 and 0.04 for AB and P_1_ respectively. That the standard deviation of the CVs stayed relatively constant across both constant Ts and T gradient conditions and orientations, while the magnitude of the CV doubled for the various T gradient conditions when compare to the constant T divisions, implies that the presence in the gradient imposes greater variability in division timing.

Second, we found that the division rates of both cells, independent of the orientation of the embryos in the gradient, decrease in the T gradient relative to their expected T-dependent behavior at their local T. This effect suggests that embryos respond to the discordant conditions by reducing the overall rate of development. Regardless of the mechanism underlying this process, many embryos showing this greatly reduced developmental rate survived, revealing their ability to adjust to these highly aberrant conditions (Fig. 3 and see below). There were two exceptions to the general trend of slowing relative to the rate expected for T environment. First, for embryos that experienced a T gradient of 5°C starting at the 1-cell stage, the division rates were more consistent with the expected behavior for the local T (Fig. 4I and J). Moreover, the rate of division of P_1_ in embryos exposed to the largest gradient (Fig. 4H) similarly deviated less dramatically from the expected behavior. In both cases, embryos in the cohorts that tracked more closely with expected timing were much more likely to die, a trend that we observed more generally as well (see below).

### On average, the warmer cell slows more than the cooler cell in viable embryos irrespective of orientation

While imposition of the T gradient slowed the division rate of both AB and P_1_, it was possible that a compensation process that normalizes the division sequence might occur in which one cell is subject to greater reduction in division rate than the other, depending on orientation in the gradient. To assess the extent to which the cell division rate was altered, we analyzed the division timing of each cell (AB and P_1_) relative to the other. For each embryo analyzed, we determined the fold change in division timing for each of the cells of the two-cell embryo by calculating the log_2_ of the ratio of observed and expected time of division at the average T experienced by each cell. The behavior of each embryo was then graphed as a single point, with the behavior of AB plotted on the x axis and P_1_ on the y axis (Fig. 5). This treatment allowed us to simultaneously identify how each cell behaved relative to both its expected behavior and to that of the other cell, as explained in Fig. 5A. The results allowed us to compare the deviations in developmental timing of each cell relative to the other in the context of the entire embryo with those measured in embryos developing at constant T. Consistent with the high fidelity of early *C. elegans* development, we found that embryos at constant T showed low variation in cell division rates around the origin of both axes [log_2_ (expected: observed) = ∼0]. If division timings of both cells slowed by the same magnitude relative their expected timings, the results would fall on a line with slope = 1, and the distance from the origin along this line would reflect the overall slow-down as a result of the gradient. Divergence from this line indicates that the division rate of one of the two cells deviated from the expected division rate by a larger extent than the other cell.

**Figure 5.**
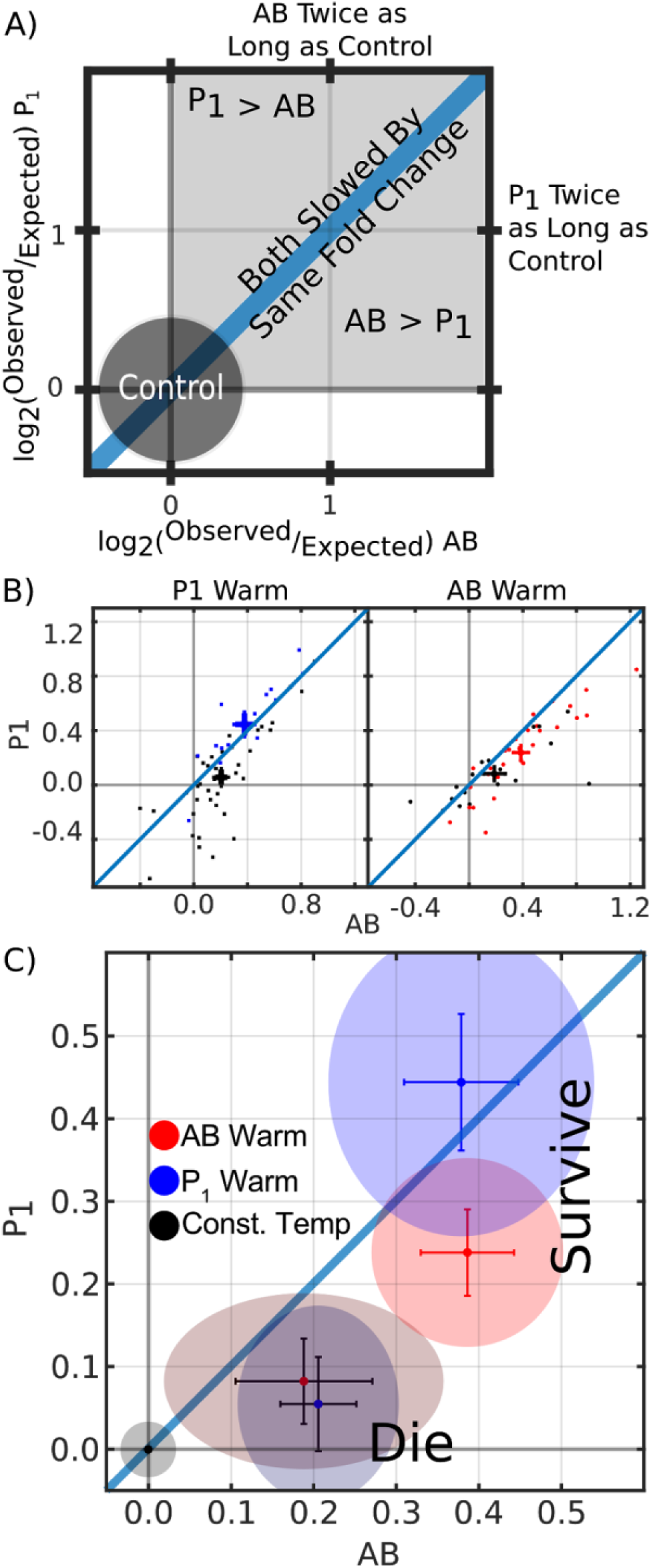
Analysis of deviations in division rates of AB and P_1_. A) Explanatory graph for fold change in division time for whole embryo. B) Scatter plot for whole embryo behavior when (left) P_1_ is warm and (right) AB is warm. Black data points represent embryos that did not survive to hatching while blue and red represent those that did. Crosses and error bars are mean and SE. C) Mean, SE, and 95% CI for fold changes in AB and P_1_, based on orientation and survival. Blue-embryos with P_1_ warmer than AB, Red-embryos with AB warmer than P_1_. Black standard error bars identify populations that did not survive. Colored ellipses represent two-dimensional 95%CI for each population. The black data point at origin and the surrounding gray ellipse are the mean and 95%CI respectively of control embryos at a uniform T.

The data were partitioned into four groups. Results for embryos in each orientation (AB warmer *vs.* P_1_ warmer) were averaged and plotted separately. Further, to assess whether the degree of deviation in cell division timing might correlate with successful embryogenesis in the discordant conditions, the results were further separated based on the ultimate outcome (viability *vs.* lethality) following exposure to the gradient.

This analysis revealed a striking outcome: embryos that developed and hatched (i.e., were viable) showed a substantially larger average overall reduction in division timing from that expected at constant T (greater distance from the origin) compared to the cohort of embryos that failed to hatch (i.e., were lethal) (Fig. 5C). Thus, survival correlated with greater slow-down in division rates of both AB and P_1_, irrespective of orientation in the gradient.

For the cohort of embryos that survived, we found that the two cells showed a pronounced orientation-dependent difference. For viable embryos positioned in the gradient with P_1_ on the warmer end, the slow-down in average division rate from the expected rate was greater for P_1_ than for AB. Similarly, viable embryos in which AB experienced the warmer environment showed a greater reduction in average division rate of AB compared to P_1_ (Fig. 5C).These results suggest that, for surviving embryos, while the division of both cells slowed dramatically in the T gradient, the cell on the warmer side tended to respond to the discordance to a greater degree relative to the T it experienced than that on the cooler side, with the outcome that normal division sequence was preserved.

We observed a second orientation-dependent effect: viable embryos in which P_1_ was warmer(P_1warm_) than AB experienced a greater magnitude in slowdown of P_1_ relative to the extent of slowdown of AB in the viable AB_warm_ embryos. Regardless, in both cases, the cell on the warmer side of the gradient always showed greater deviation from the expected rate. AB, the faster dividing cell under normal conditions, appears to be more effective at responding to the discordance than P_1_, the normally slower developing cell, a finding that is consistent with the orientation-dependent effect on lethality described above (Fig. 5C).

In contrast to the results with viable embryos, the cohorts of inviable embryos tended to show a substantially reduced average response of both AB and P_1_: in both orientations, the data clustered closer to the origin than for the viable embryos. Moreover, unlike the surviving embryos, these inviable embryos showed a greater reduction in division rate of AB compared to P_1_ in *both* orientations, with the result that the data for the two orientations clustered together (Fig. 5C).

In summary, we found that both AB and P_1_ greatly reduce their division rates in embryos that survive the T gradient and that the cell that would be expected to divide more rapidly on the basis of its higher T environment shows a larger response (greater reduction in division rate) than its cooler neighbor.

### Evidence for a later compensation mechanism: morphogenesis can progress despite reversal in the AB and P_1_ division sequence

Under the most extreme conditions, we found that the T gradient was sufficient to force reversal of the stereotyped division sequence of AB and P_1_ (Fig. 6). For embryos subjected to the gradient after cleavage of P0 and oriented with P_1_ on the warmer side, we found that a steep T gradient was sufficient to reverse the normal division sequence and drive P_1_ to divide before AB in 60% (9/15) of embryos subjected to a 6.5°C gradient, and 71% (12/15) of those experiencing a 7°C gradient. Further, initiation of the gradient prior to the division of P_0_, in which the posterior side of the embryo was oriented toward the warmer end of gradient, resulted in reversal of the division sequence in 90% (9/10) of the embryos. As expected, none of these embryos in which the sequence of division was reversed survived and hatched. Unexpectedly, however, a substantial fraction of such embryos proceeded through relatively normal morphogenesis: 32% (9/28) of embryos that experienced a reversal in division sequence of AB and P_1_ gave rise to an arrested embryo that appeared relatively normal in morphology and had undergone substantially normal morphogenesis (Fig. 6). Moreover, we found that 43% (6/14) of lethal embryos that experienced any of the gradients in which AB was warmer than P_1_, similarly proceeded through relatively normal morphogenesis. These findings revealed that even under extremely discordant conditions that drive complete reversal of the stereotyped division sequence in the very early embryo, later embryos appear to be capable of compensating sufficiently well that morphogenesis, if not fully successful development, can occur. These observations underscore the substantial ability of *C. elegans* to correct for aberrations in cell division and placement patterns.

**Figure 6.**
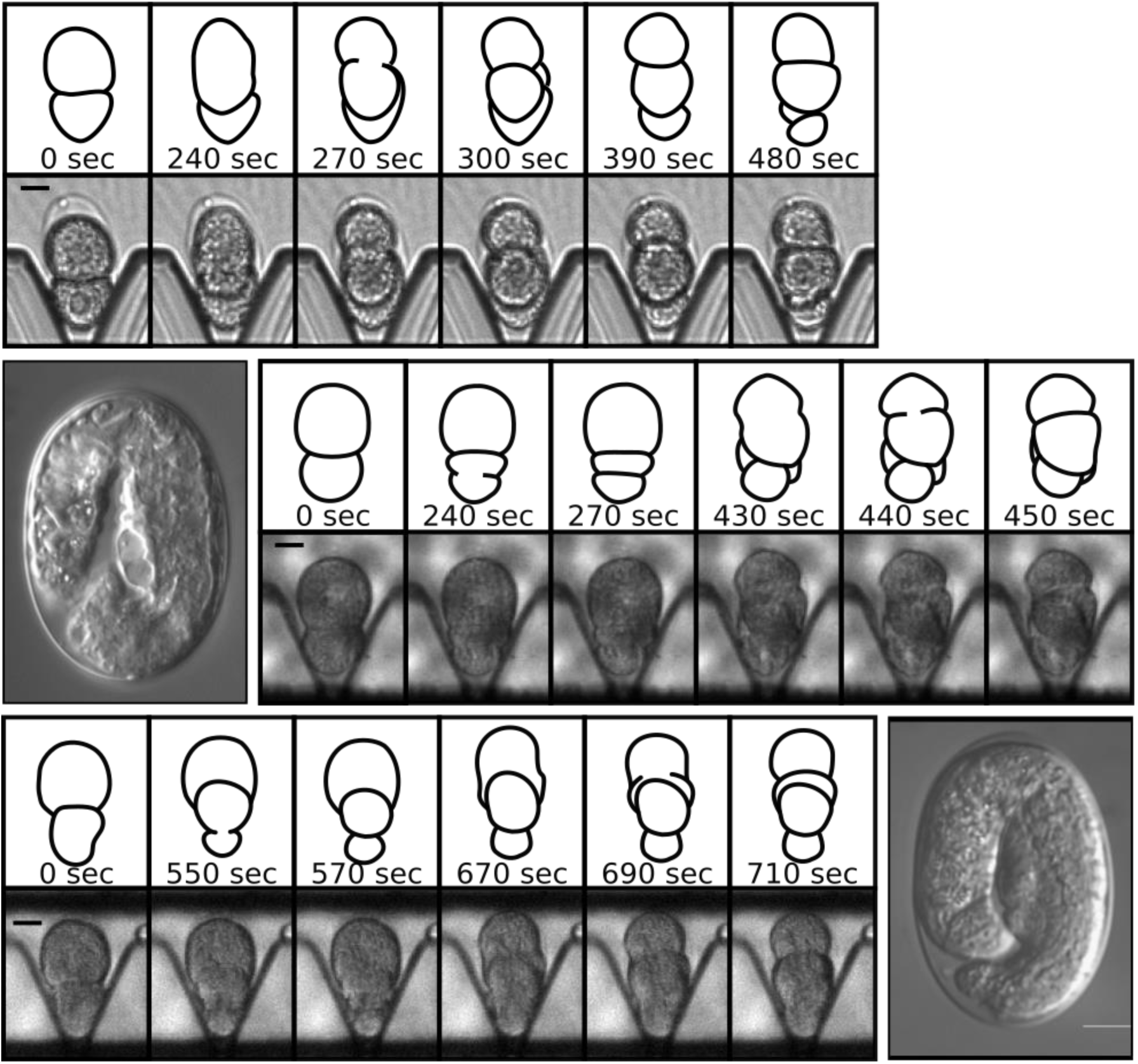
Relatively normal morphogenesis following out of sequence divisions of AB and P_1_. Time lapse images and outline of early cell divisions of AB and P_1_. Top panel: stereotyped control embryos with the larger anterior AB cell dividing before the smaller posterior cell P_1_ at a uniform permissive T. Bottom and middle panels: example of two embryos experiencing a reversal of the division sequence of AB and P_1_, along with 100X DIC image of an arrested embryo∼ 24 hours after being in T gradient. The embryo had progressed through morphogenesis and elongation despite the reversed sequence of AB-P_1_ divisions.

## Discussion

A largely unexplored problem in animal biology is how complex developmental processes result in a reliable output in spite of constant environmental and intrinsic noise. Our goal in this study was to test whether animal embryos that normally show a highly stereotyped pattern of development are capable of responding to, and correcting for, the discordant conditions imposed by a T gradient. We propose that this discordance is a proxy for natural noise that embryos normally experience which, in the absence of any correction, might otherwise cause them to deviate from the stereotypical pattern. These studies demonstrated that *C. elegans* embryos both respond to a T gradient and can adjust division timings to generate a normal pattern of development despite highly discordant T’s.

In this study, we report the following major findings. 1) We have designed, fabricated, and validated a microfluidics device that effectively establishes a steep T gradient of up to 7.5°C across the 50 μm anteroposterior axis of *C. elegans* embryos. 2) We have characterized the division time of the two-cell embryo as a function of T and established a mathematical model describing the relationship. 3) Embryos through at least the second round of division can survive exposure to a pole-to-pole gradient of up to 7°C and hatch into viable larvae. 4) Survival in the T gradient is orientation-dependent: embryos in which AB is positioned on the warmer side can withstand a larger gradient than those in the reverse orientation. 5) Embryos exposed to the T gradient slow their average developmental rate dramatically compared to those embryos that do not survive to hatching at the same gradient magnitude. 6) While the rate of division of both cells at the two-cell stage is reduced in the gradient, the cell on the warmer end shows a tendency to slow by a larger degree than that on the cooler end, thereby often preserving the normal geometry of the embryo. 7) Survival correlates with the magnitude of this cell division response: the cohort of embryos that died showed a lower average deviation in division timing from that predicted based on their T environment. 8) Some embryos in which the AB/P_1_ division sequence was reversed invariably died but nonetheless showed signs of relatively normal morphogenesis, suggesting the existence of later developmental compensation mechanisms.

### Evidence for multiple compensation/correction systems in embryos

Mechanisms that detect and correct for “errors” resulting from noisy development processes might function by (A) sensing deviations in rates or timing of events outside normal bounds and adjusting for these deviations at the time of their occurrence, or (B) by acting at pre-established stages (perhaps “checkpoints”) to detect aberrant events that have occurred in the past, and make repairs through subsequent compensation processes. Our findings are consistent with both types of systems in *C. elegans* embryos.

The finding that the division of both cells slows in the gradient relative to the expected behavior is consistent with the possibility that the discrepancy from normal development imposed by the gradient activates a checkpoint-like system in which a “crisis” leads to slowing or pausing of the cell cycle (as occurs, for example, in genotoxic-induced stress [29,53–55]. This effect correlates with survival: the cohort of embryos that survived showed the most dramatic reduction in developmental rate. We postulate two possible explanations for this effect. First, it is conceivable that T gradient across each cell induces a cell-intrinsic process that slows the rate of division in response to the aberrant environment independently of effects on the other cell. A second possibility is that such a response might result from intercellular communication between AB and P_1_ that instructs both cells to slow or “wait” until adjustments to division rates have been made. In such an event, this cell-extrinsic communication would be bidirectional, as both cells slow down relative to their expected behaviors. Thus, each cell at this, and possible later stages, might compare its progress with neighboring cells and “tune” its division timing in such a way that the proper geometry is ensured. Resolution of these alternatives would require creating a step-gradient in which two sharply delineated T’s are imposed upon the cells, where each T is experienced uniformly across the full dimension of the cell. If the effect we have observed also occurs under such conditions, it would strongly argue that the effect is mediated through cell-extrinsic signaling.

We found that as early as the two-cell stage, embryos show evidence of “tuning” of cell divisions in response to deviations: under conditions in which each cell would be expected to divide at an inappropriate time relative to the other, AB and P_1_ often appear to adjust their division rates in a way that maintains their normal relative division sequence. It is therefore conceivable that a cell that might be driven to divide more rapidly as a result of a warmer T might alter its division rate based on information about the rate of its cooler neighbor. Our results suggest that when its cooler neighbor is lagging, AB shows a greater capability for slowing its division than does P_1_. This is reflected both in the magnitude of the division rate decrease (Fig. 5) and the higher viability of embryos in which AB is located on the warm side than those in reverse orientation. Given that AB is the “leader” during normal development (i.e., it divides before P_1_ under constant T conditions), this may reflect the intrinsic ability of AB under normal conditions to monitor and respond to its slower neighbor as needed to maintain the proper relative division timing. Regardless, if AB and P_1_ undergo intercellular communication to regulate developmental progress, it could explain the observation that isolated AB blastomeres, obtained by removal of P_1_ by extrusion from the eggshell or through isolation in culture, undergo slower rates of division ([27,56,57]; our unpublished observations); in the absence of information from P_1_ that might indicate progress in its development, AB may default to a slower division rate. Our findings also raise the question of whether the adjustment in cell division timing observed here is related to a different cell timing compensation mechanism: the negative correlation between cycle timing of a cell and its descendant, in which cells that divide early give rise to granddaughters that are more likely to divide late [46].

Our observations suggest that the capacity of embryos to compensate for the discordance of the T gradient can be exceeded beyond an acceptable “dynamic range” under extreme conditions. Most or all embryos fail to complete normal embryogenesis when exposed to the largest gradient (Figs. 3 and 5). We note that under these extreme conditions, the magnitude of the overall slowdown relative to the expected rate is less than under milder T gradient conditions, suggesting that the response system may be overwhelmed by this environment. It is also striking that the cohort of surviving embryos show the greatest average reduction in both overall cell division rate and in relative slowdown of the warmer vs. colder cells: in the viable embryos the warmer cell division rate slowed down related to its expected rate by a larger factor than that of the cooler cell, hence restoring what would have otherwise been an out-of-sequence division pattern. Thus, it appears that the embryos that respond most dramatically and correct the discordance most effectively are the most likely to survive. We propose that this effect may reflect an active process that corrects for noise-induced drift and ensures a faithful output.

Our data also support the existence of a later-acting compensatory system that correct errors after the early cell divisions. Although some “P_1_-warm” embryos failed to correct for the discordant conditions and reversed the stereotyped division sequence, they could nonetheless proceed through apparently relatively normal development, resulting in a worm-like, albeit lethal, animal (Fig. 6). Rapid signaling events in the early embryo depend on precise geometry of cells (for example, in the induction of both gut in the EMS lineage and of ABp-specific fate by the P_2_ cell; [22,58–64]). Our observations suggest that morphogenesis can be coordinated and corrections made even after an embryo with aberrant cellular arrangements has formed. This finding is consistent with reports that, at elevated T’s, much later mid-stage embryos show variability in cell positions and cell lineage patterns and yet resolve into normal healthy animals through normal cell repositioning and morphogenesis [21,65,66].

### Potential regulatory processes in compensation to discordance

It will be of interest to understand the molecular machinery that might mediate the profound cell division timing adjustments we have observed in embryos exposed to the T gradient. The asynchrony of AB and P_1_ division timing is known to reflect at least two checkpoint-based regulatory systems. First, cell-size-dependent control by ATL-1 and CHK-1 accounts for approximately 40% of the difference in cell cycle timing between AB and P_1_ [29]. The other system acts independently of cell size and depends on localization of PLK-1 and CDC-25.1 in P_1_ [67,68]. The tight regulation of the cell cycle seen in early in *C. elegans* embryogenesis is also apparent in mice, in which DNA damage and spindle assembly checkpoints are active [69,70]; however, this does not appear to be the case in other vertebrates, including Xenopus and zebrafish [71,72], in which these cell division regulatory systems are enabled only after the midblastula transition. It is conceivable that regulatory events that influence either the ATL-1/CHK-1 checkpoint system or the PLK-1 checkpoint system in the early *C. elegans* embryo could mediate the response to discordant conditions of the T gradient.

A prominent example of a system that coordinates cellular timing during development, thereby ensuring highly reproducible pattering, is the segmentation clock for somitogenesis in vertebrate embryos, which is controlled in part by Notch signaling [17–19,73]. Notch signaling is also used to specify cell identities throughout development in *C. elegans*, including in the very early embryo [22,74]. It is intriguing to note that the maternally provided GLP-1 Notch receptor is differentially expressed as early as the two-cell stage in *C. elegans*, where it is translated in AB but not P_1_. Moreover, LIN-12, the other Notch receptor, is zygotically expressed as early as the 24-cell stage [75] and its (presumably maternal) transcript has been detected at low levels as early as the one-cell stage [76,77]. While it is tempting to speculate that Notch signaling might function in coordination of AB and P_1_ division, no Notch ligand has been found to be expressed as early as the two-cell stage, though the APX-1 ligand is expressed, and functions, in its daughter P_2_; [63]. It will be of interest to test whether early embryos lacking both Notch receptors show an altered response to a T gradient, in which case Notch signaling might be implicated in mediating communication between AB and P_1_ that coordinates their division timing.

### Other potential implications of compensation to a T gradient

Our findings may be relevant to understanding the factors the dictate the T limits over which poikilotherms are able to develop successfully. It is generally accepted that these limits reflect, in part, the T range over which critical intracellular components are able to function properly. Studies of closely related species of nematodes and flies demonstrate that there is a uniform scaling of development in time as a function of T [51,78]. The exponential nature of the T-dependent rates of events in the one-cell *C. elegans* embryo [51], and the T-dependent model of the first cell divisions described here, raise the possibility that the failure of development at T’s that are just outside the range for successful development might also be the result of divergence of cell division clocks within the developing animal as a result of a breakdown in the compensation system.

Finally, the apparent compensatory system suggested by these findings may account for observations that human embryos are particularly susceptible to failure early during embryogenesis, which appears to be coupled with mismatches in cellular timing in very early embryonic cells [79]. An early-acting system that detects and compensates for cell-cycle timing defects might function as a global developmental abort system in higher organisms, particularly when full-term development is costly.

## Materials and Methods

### Construction of a microfluidic device to generate a stable steep T gradient

The microfluidic device consists of two main layers: a backplane, containing the vias and electrodes of the device and a second layer of microchannels placed on top of the backplane (see supplemental text for detailed method). The device uses a platinum Joule micro-heater to establish the high T side of the gradient. The Joule heater along with four micro resistive thermal devices (RTDs) acting as local T sensors, are simultaneously patterned through micro-lithography and metal deposition on a glass substrate. Electrical current generates an approximate cylindrical dispersal of heat around and away from the Joule heater. To reject the heat and focus the T gradient, a chilled fluid mixture is flowed underneath and in contact with the glass substrate on the surface opposite from that containing the heater. The magnitude of the T gradient as well as its rate of change is controlled by varying the T of the fluid underneath the glass, as well as the power through the Joule heater. A microfluidic channel with “trapping pillars” to capture and orient *a C. elegans* embryo within the T gradient, is placed on top of the heater. The microfluidic channel terminates at the end of the glass substrate where microbore tubing is affixed in a manner that allows embryos to be introduced into the device.

### Characterization of the in-device T gradient

The T gradients generated in the microfluidic device were characterized using thermometric microscopy to correlate the fluorescence intensity of Rhodamine B with T and measurements from on board resistive thermal sensors or devices (RTDs). After ensuring that there are no bubbles in the microchannels, the water was replaced by a dilute solution of dextran conjugated Rhodamine B (DCRB). To construct a standard curve relating T at each point in the channels with fluorescence intensity, approximately 30-60 fluorescence images of the DCRB filled microchannels are taken at each T between 30°C and 1.5°C which is achieved by regulating the voltage applied to the Joule heater. The T at each point in the device is estimated by the classical least squares model for each pixel. 2D Finite element analysis was performed using Comsol Multiphysics versions 5.1-5.2a. Built-in material properties were used, with the notable exceptions of the physical parameters for SU-8, and NOA 81, which were both estimated to have the physical properties of polyethylene. Cooling fluid flow under the device was assumed to be laminar. The resistance measured by the different RTDs in the gradient was correlated with the T using standard least squares fitting (see supplemental text).

### Embryo preparation and loading

Embryo experiments were conducted with the *C. elegans* laboratory reference strain N2 which was maintained as described by Stiernagle [80]. Strains were maintained at either room T (18-22°C) or in a 15°C incubator. Young adult worms are cut open under a dissecting scope in osmotically balanced Edgars egg salts solution as described by Edgar and McGhee [59] and selected embryos are transferred via mouth pipette to the end of tubing that is connected to the inlet ports of the microfluidic device. One-cell embryos were loaded into the device either before pronuclear meeting or immediately after. For two-cell embryos, we continued to track development outside of the microfluidic device until the first membrane cleavage, at which point they were loaded into the device, using a syringe pump. Embryos generally reached the capture region of the device within 30-60 seconds. A flow of 500nl/min was maintained in the device while embryos were in the device to ensure that they did not experience hypoxic conditions. Embryos were unloaded from the device by operating the syringe in reverse. The embryos were then transferred with a mouth pipette to a standard agar plate seeded with E. coli OP50 and incubated at room T and scored if they hatched or were dead ∼ 24 hours later. Embryos with reversed sequence of cell division were observed 24 hours later on a Nikon Eclipse Ti at 100X magnification

### Estimation of cell division rate

Embryos were imaged on an upright Nikon Microphot microscope at 10 second intervals. Cell division interval was determined as time between successive cytokinesis as inferred by the first image that shows apparent completion of membrane pinching. For embryos loaded after the first division, the rate of division was estimated using the measured room T and the linear model of time of division as a function of T.

## Acknowledgements

Nematode strains used in this work were provided by the Caenorhabditis Genetics Center, which is funded by the National Institutes of Health - Office of Research Infrastructure Programs (P40 OD010440). This work was supported by grants from NIH (#1 R21 HD075292).

## Supplementary Text

### Device construction

The microfluidic device is constructed as two main layers: the backplane, containing the vias and electrodes of the device and a second layer of microchannels placed on top of the backplane. Three input and output tapering hemi-conical vias, approximately 2 mm wide, and 800-900 μm deep at the edge, are made in a 1”x3” commercially available microscope slide cut in half to 1”x1.5”. To prevent impurity migration from the microscope slide, 100-150 nm of SiO2 is reactive sputter deposited on the backplane. Electrodes of 10 nm Ti followed by 100 nm of Pt were patterned on microscope slides using standard negative photoresist clean room photolithography. The electrode face of the device is covered with an approximate 2 μm layer of SU-8 2002 (MicroChem Corp., Westborough, MA). Centered on the device, an ∼ 1cm diameter and 750-800 μm deep circular cut, under and in the opposite face of the glass, is HF etched into the glass which allowed a steep T gradient of 7.5°C by reducing the thickness of the coverslip and ensuring the heat was dissipated away by circulation of chilled water. A positive master mold of our microfluidic channel design was dry etched 40 μm deep into a 3” silicon wafer with a negative PDMS mold made from the silicon wafer using standard soft lithography techniques [1]. The microfluidic “sticker” layer of the device is constructed consistent with methods developed by Bartolo et al. [2], and is placed on top of the electrodes with the capture pillars of the device centered on the electrodes of the backplane. 0.03” outer diameter PTFE tubing is inserted into the device vias, and secured with two-part epoxy.

Device is mounted on a custom-built device holder/flow cell that allows bulk fluid flow underneath and in contact with the outside bottom surface of the microfluidic device. Tubing (∼0.25” ID PDMS) connects the flow cell to a water circulator filled with DI water and ethylene glycol in a ratio of 4:1. Flow rate through the fluid cell is on the order of 19 ml/sec. The water circulator is used to set the background T of the device. Device holder/flow cell and device are loaded into a custom rig on an upright microscope. A custom environmental chamber enclosing the microscope is maintained at a slight positive pressure with sub 0°C dew point laboratory supplied air to prevent condensation on device during operation.

### Characterization of the T gradient

Effect of T on the fluorescence response of Rhodamine B has been extensively studied and its quantum yield is highly T dependent[3–5]. It has been previously reported that a solution of dextran-conjugated fluorophores can aggregate, resulting in an apparent increase in quantum efficiency of the fluorophore, and that the aggregation rate is T dependent [6]. To address this concern, we performed our measurements with a flow of the solution running during measurements. To ensure the introduced flow would not affect the T profile of the device we calculated the expected flow rate of our device for which the Pe would equal one and found it to be on the order of 1-2μl/min. We then measured the effect of fluid flow above and below this threshold with thermometric microscopy utilizing DCRB. We found that a flow of 2μl/min did not affect the T profile of the gradient, while at flow of 15μl/min shifted the T profile in the direction of flow. In later devices, in addition to characterization of the T gradient with thermometric microscopy, we also included resistive thermal sensors or devices (RTDs) [7] in the T gradient region of the device. The device is placed in a well stirred ice bath and allowed to come up to room T while measuring the T of the bath with thermocouples and resistance of the RTDs. Standard least squares fitting is used to relate the RTD measurements with T. We found a highly linear correlation between T and the measured resistance, and modeled the relationship between the two using a least squares linear model. R^2^ values of linear models fitting resistance to Ts ranging from 0°C to 20°C were typically on the order of 0.999. To verify that the RTDs were primarily measuring the T in the region of the T gradient, and not the electrode leads leading up to that region, we measured the resistance of the patterned RTDs with the device mounted on a flow cell that flowed a fixed and measured T of water below the region of the device where the T gradient is established. We found that our T measurements were within 1°C of the experiment in which the device was fully submerged.

During normal operation of the device, the device is not submerged in fluid. To verify that our T measurement was a reasonable estimate of the T in the channel, and not just the bottom of the channel, we constructed a modified flow cell that allowed the flow of the T setting water both underneath as well as over the top of the device. We found that the average difference in T measured between when the top of the device is exposed to air, and when it is sandwiched between flowing fluid was on the order of a third of a degree.

### Modeling the apparent intra-embryonic T gradient

We modeled the embryo as a 50×30 μm spheroid with thermal conductivity equal to that of water, k_cytoplasm_=0.6 W/m-K [8], and an insulating eggshell of 300 nm thickness [9]. Although cytoplasm is a gel matrix, thermal conductivity of a gel, for example a concentrated protein solution of 10% gelatin, is only 5% lower in conductivity than water [10]. While thermal conductivity of nematode eggshells has not been measured, a model of *Drosophila* embryos [8] used k_shell_=k_paraffin_ wax =0.25 W/m-K, 10x more insulating than an avian eggshell. Using this extreme value in our simulation, the intra-embryonic gradient was reduced by only 1%. We also considered the possibility that extremely active fluid circulation within the embryo might overcome the T gradient within the embryo by convective transport. The Peclet number (Pe) of a system indicates whether convection or diffusion dominates in determining the distribution of heat. A Pe of one indicates a system where convection and diffusion are in balance. Values higher than one indicate convection dominates and values lower than one indicate diffusion dominates. The maximum known cytoplasmic streaming velocity in the *C. elegans* embryo of 7 μm/min [11], cannot overcome thermal diffusion at this scale as the Peclet (Pe) number of the embryo with known dimensions and expected possible highest velocity is only 2.5×10^−5^. Thus, the T gradient within the embryo is in close accordance with the external T gradient in the microfluidic device.

### Verification that the microfluidic device is compatible with embryonic development

To verify compatibility of the device with embryo development, a cohort of early stage embryos (1-8 cell stage) were loaded into the device and allowed to develop to hatching while in the device. Below a certain threshold of flow, the embryos tended to arrest during development and or not complete development. This finding was consistent with the material from which the device was constructed, NOA81, being gas impermeable [12]. Flow rates in excess of 25nl/min prevented arrest of embryos during development. Having previously calculated the Pe of the device and measured the effect of fluid flow below the critical rate, we were confident that a flow rate of 100-500nl/min within the device would not affect the T profile of the device while simultaneously creating a biologically compatible environment. Our real time RTD measurements of T in the device in our later experiments also demonstrated that the T profile at these slower flow rates remained similar to those without flow. To determine the effect of loading and unloading on the survival of the embryos, we loaded a cohort of one-celled and two-celled embryos into a room T device at 80μl/min, left them in the device for ∼ 1 hour, with a trickling flow of 500nl/min and unloaded them at a rate of 300μl/min. Each embryo was then placed on an agar plate and evaluated for whether or not they had successfully developed and hatched 24 hours later. We found that the rate of hatching was 98.4% (61/62). We were thus able to optimize the parameters that ensured the viability of the embryos was not adversely affected in the microfluidic device under control conditions of uniform T.

## Supplementary movies

**Movie S1**

Animation of fly-over and through of device. All device sizes are approximately to scale relative to the bulk substrate of the device which has an aspect ratio of 1:3:1/25.4 (aspect ratio of a typical commercially available microscope slide)

**Movie S2**

High speed camera acquisition of loading of a single embryo were taken at 10k frames per second at 10x on inverted Nikon Eclipse. Replay speed was 10 frames per second.

## Notes

### Competing Interest Statement

The authors have declared no competing interest.

